# A control policy can be adapted to task demands during both motor execution and motor planning

**DOI:** 10.1101/2023.10.16.562495

**Authors:** Jean-Jacques Orban de Xivry, Robert Hardwick

**Author notes:** Corresponding author : Jean-Jacques Orban de Xivry, Movement Control & Neuroplasticity Research Group, Tervuursevest 101 - box 1501, 3001 Leuven.

## Abstract

Movement planning consists of several processes related to the preparation of a movement such as decision making, target selection, application of task demands, action selection and specification of movement kinematics. These numerous processes are reflected in the reaction time, which is the time that it takes to start executing the movement. However, not all the processes that lead to motor planning increase reaction time. In this paper, we wanted to test whether tuning the control policy to task demands contributes to reaction time. Taking into account that the tuning of the control policy differs for narrow and wide targets, we used a timed response paradigm in order to track the amount of time needed to tune the control policy appropriately to task demands. We discovered that it does not take any time during motor planning and even that it can occur indistinguishably during motor planning or during motor execution. That is, the tuning the control policy was equally good when the narrow or wide target was displayed before than when it was displayed after the start of the movement. These results suggest that the frontier between motor planning and execution is not as clear cut as it is often depicted.

**New & Noteworthy:** Movement preparation consists of different processes such as target selection, and movement parameters selection. We investigate the time that it takes to tune movement parameters to task demands. We found that the brain does this instantaneously and that this can even happen during movement. Therefore, this suggests that there exists an overlap during movement planning and execution.

## Introduction

Movement preparation involves the integration of multiple sources of sensory information to allow the execution of coordinated and efficient actions, and has been conceptualized as a two-stage process (1). The first stage involves a non-motor perceptual “decision making” process through which the goal of the movement is selected, involving the sub-processes of detecting relevant stimuli and identifying the target of the action, both of which can be modified by relevant task constraints. This is followed by a second “motor planning” phase wherein the action to execute is selected, relevant movement kinematics are applied, and the final movement to be executed is specified via a motor command. Classical studies indicate that the time required for movement preparation increases with the complexity of the task demands (2). In line with this suggestion, contemporary work has proposed that the majority of the time required for movement preparation is in fact consumed by the decision-making process, arguing that motor planning itself can be achieved almost instantaneously (1). This is, however, at odds with the finding that adapting the control policy to task demands carries a switching cost (3, 4). A switching cost refers to the fact that changing the task demands decreases performance, and that participants behave differently immediately after a change in tasks than when the same task demands are repeated for several trials in a row (3, 4). Such switching costs typically influence the reaction time (5).

The motor planning phase of movement preparation is often examined within the framework of optimal control theory (6, 7). Optimal control theory proposes that movement selection minimizes a cost function that balances a tradeoff between movement accuracy and energy expenditure. This has widely been studied by examining whether movements respect the ‘minimum intervention principle’ in the presence of perturbations or of natural movement variability (8). If a perturbation does not interfere with the goal of the movement (i.e. it introduces ‘task-irrelevant variability’), the response will be relatively minor; by contrast, perturbations that interfere with the goal of the movement (i.e. that introduce ‘task-relevant variability’) will be met with more robust corrections (9– 11). The absence of correction for task-irrelevant dimensions has also been linked to an uncontrolled manifold (12–14, but see 15). The uncontrolled manifold corresponds to a space of movement parameters that are irrelevant to task success and that are left uncontrolled by the motor system. When reaching to a narrow target, both optimal control theory and the uncontrolled manifold hypothesis would predict that any perturbation that affects hand trajectory and impedes task success should be strongly resisted. In contrast, if the target is wide and the perturbation does not reduce task success, the response to the perturbation should be much smaller, as found in numerous studies (3, 4, 16–19). The minimum intervention principle extends to error correction mechanisms as well, as errors are only corrected in task-relevant dimensions (3, 4, 20). In the context of the optimal feedback control theory, the reaction to a perturbation or to natural movement variability is determined by the control policy, which is presumably determined during the motor planning stage, hence during the reaction time period (1).

Using a postural perturbation paradigm, Yang and colleagues (33) investigated the time that was necessary to integrate one of two instructions into the feedback response to a perturbation during posture and found that it took 90ms for the brain to integrate the right instruction into the long-latency stretch response. However, it is unclear whether modulating task instructions is the same as adapting the cost function with target width. In addition, we want to investigate the influence of the cost function during movement while Yang et al. investigated postural movements. For movement planning, the computation occurring during the reaction time period can be assayed via timed response paradigms that limit response preparation time (21). Using such an approach, it has been demonstrated that movement preparation and initiation are two separate processes (22). This paradigm allows one to force participants to initiate their movement before movement planning is fully completed (22, 23).

Here we used the timed response paradigm to investigate how fast the control policy can be tuned to task demands, i.e. to their ability to implement the minimum intervention principle (7). If the time required for computing the optimal control policy is truly negligible (1), then limiting the time participants have to prepare movements should have a discrete, stepwise effect on the adjustment of the control policy to task demands (i.e. only movements prepared with sufficient time would show changes in response to task demands, such as responses to perturbations). By contrast, if the time available to prepare movements does influence the tuning of the control policy in function of task demands, we could instead expect that changes in the control policy in response to task demands would gradually increase with the amount of time available.

## Materials and Methods

### Participants

Forty healthy subjects (29 female, 11 male, mean age±SD = 21.1±2.0 years) participated to this experiment. Participants were right-handed according to the Edinburgh Handedness Inventory (24), and free from neurological dysfunctions. All participants provided written informed consent prior to the experiment. All procedures were approved by SMEC (Sociaal-Maatschappelijk Ethische Commissie) of the KU Leuven (G-2019 02 1536).

### Setup

Participants sat in front of a robotic arm (Endpoint Kinarm; Kinarm, Kingston, ON, Canada), and held the right handle with their right hand. A force transducer was placed below the handle in order to record the force applied by the hand against the robot. The participant’s own right hand was hidden from view by a mirror that reflected the image of a monitor placed above the mirror. The distance from the hand to the mirror was equal to the distance between the mirror and the display, making the cursor appear in the same position as the participant’s hand. The computer that controlled the robotic manipulandum stored hand position, velocity, and force at 1000Hz for later offline analyses.

## Protocol

### Forced preparation time condition

In a forced Preparation Time trial (Fig.1A), participants first moved the white hand cursor within the starting position (red square of 0.36cm^2^). After a time period of 400ms, four consecutive tones were played every 333ms for a total time interval of 1s. Participants were instructed to start of their movement synchronously with the fourth tone. The target always appeared at the same location (centered 15cm above the start position) at one of the ten possible instructed preparation times: between 594 and 1ms before the last tone, by steps of 66ms. The target consisted of an aperture within a white circle (Fig.1B, radius of 15cm, centered on the starting position). The size of the aperture could be either narrow (1cm) or wide (8cm), and this target size varied randomly from trial-to-trial (50% of trials within each block presenting each target size). The participants were instructed to bring the cursor outside of the circle via the aperture and to stop their movements when they were outside of the circle. During the movement, the small cursor disappeared and was replaced by a white circle that was centered on the starting position, and whose radius was equal to the distance between the hand and the starting position. This expanding circle provided thus information about the extent of the ongoing movement, but not about its direction.

**Fig. 1:**
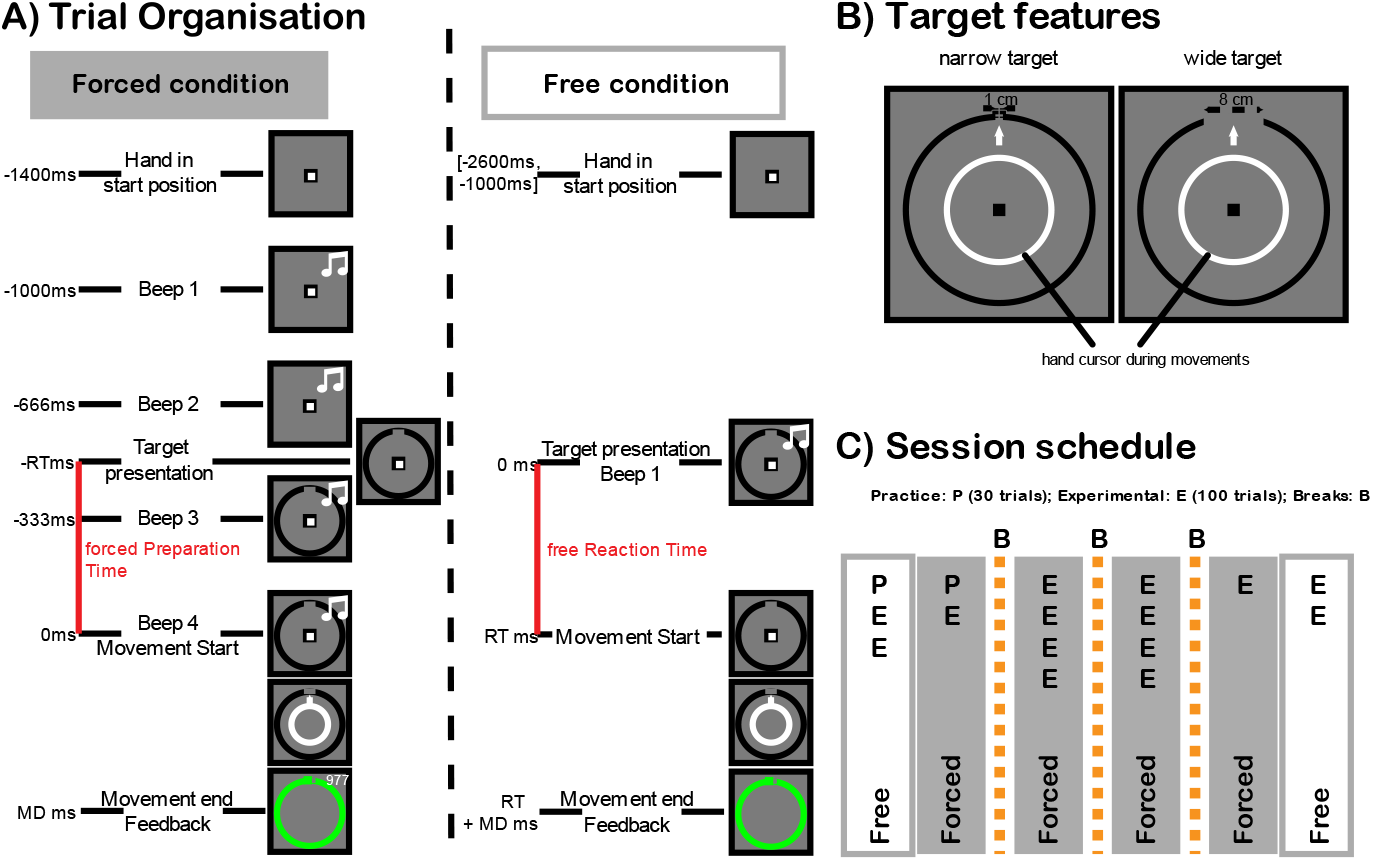
A) Trial organization: 1. Forced Preparation Time trials (Forced condition). The participants had to first bring their hand cursor (white square) inside the starting position (red square of 0.6×0.6 cm, represented by the black square). After having remained inside the start position for 400ms, a series of four beeps were delivered at a 333ms interval for a total duration of 1s. The target appeared in one of the ten possible forced Preparation Time during this interval. Subjects were trained to start their movement at the time of the fourth beep. Their movement ended when they travelled 15cm away from the start position in any direction. At that location, an imprint was presented to provide participants with feedback about the accuracy of their movement. If the imprint was within the target region delimited by the circle, this circle became green. The color of the imprint indicated to the participant whether they moved too fast (yellow), too slow (blue) or at the right speed (green). Finally, feedback about the timing of the start of the movement with respect to the first beep (in ms) was presented to the participant just above the target region. A perfect timing corresponded to 1000ms. **2. Free Reaction Time trials (Free condition):** After placing their hand cursor within the starting position, participants waited between 1000 to 2600ms for the display of the target. The target presentation was accompanied by a beep. In response to this event, participants had to elicit a reaching movement to the target. As in the forced Preparation Time trials, they then received feedback on movement accuracy and movement speed, but not on reaction time. **B) Target features:** The target consisted of a white circle (radius of 15cm) with a gap that was either narrow (1.0cm) or wide (8.0cm). The order of target presentation was randomly interspersed within each block, with 50% of trials being made to each target size. The center location of the target remained identical for all trials. During the reaching movement, the cursor disappeared and was replaced by an expanding circle that was centered on the start position in order to provide feedback on the distance (extent) of the movement, but not its direction. **C) session schedule:** Participants first performed one practice block of free Reaction Time trials followed by two blocks of 100 free Reaction Time trials. Then then performed 30 practice trials of forced Preparation Time trials followed by one block of 100 forced Preparation Time trials. After a break, four more blocks of 100 forced Preparation Time trials were performed, followed by another break, and another four blocks of forced Preparation Time trials. After the last break, one last block of forced Preparation Time trials was performed and two blocks of free Reaction Time trials.

Crossing the 15cm circle corresponded to the end of the movement. At that time, the location of the hand (i.e. the position of the unseen cursor) was recorded and displayed as an imprint (small circle of radius of 1mm), <u>providing feedback on the endpoint for that trial</u>. In the forced Preparation Time trials, three pieces of information were provided as feedback at the end of the trial. The color of the imprint indicated whether the movement was too fast (shorter than 300ms, yellow imprint), too slow (longer than 500ms, blue imprint) or correct (green imprint). The location of the cursor provided feedback about the accuracy of the movement. If the cursor was within the aperture, the target circle turned green. The time interval in milliseconds between the first beep and the start of the movement was also displayed right above the aperture in order to provide feedback about the timing of the movement with respect to the first beep. A timing of 1000ms (1ms after the fourth beep) would be considered perfect. Additional verbal feedback was given by the examiner to start earlier or later depending on whether they were ± 75 ms from this target time. Participants earned one point when the movement was accurate and one point when the movement had the right speed.

### Free reaction time condition

Free Reaction Time trials are a simplified version of the forced Preparation Time trials. In these trials, once the participant had bought their hand into the starting position, the target appeared together with a beep after a delay of between 1000 and 2600ms. Participants were instructed to start moving as soon as possible after the appearance of the target. Once the movement started, the free Reaction Time trials were identical to the forced Preparation Time trials except that no feedback about their reaction time was provided.

### Perturbation trials

A subset of forced Preparation Time and free Reaction Time trials consisted of perturbation trials. During these trials, the hand was deviated from the center of the target by virtual walls that were created by applying a stiff unidimensional spring (spring stiffness: 1000 N/m; viscosity: 10 N·s/m). The two walls were not separated by any distance. Therefore, the robot acted as a mechanical guide that directed the hand 2 cm to either the left or the right of the center of the target and forced the hand to a straight line. The perturbation started when the hand was still in the starting target and lasted until the hand travelled 15cm (and exited the circle). At the end of these trials, the imprint was always displayed in the center of the target to hide the fact that a perturbation was applied. Given the use of the expanding circle during the movement, no visual information about the perturbation was provided to the participants.

### Session schedule

The session was divided into practice and experimental blocks (Fig.1C). Two practice blocks gave participants the opportunity to familiarize themselves with the task, and to make sure that the instructions were clearly understood. Practice blocks were 30 trials long and contained no perturbation trials. Data from these blocks was not analyzed.

Experimental blocks of free Reaction Time and forced Preparation Time conditions contained 100 trials, 20% of which included perturbations. To ensure that we had an equal number of perturbation trials per target size (2 levels) and instructed preparation times (10 levels), we created 10 different forced Preparation Time blocks that contained a defined trial order. The order of these 10 blocks was then randomized for the different participants. These 10 blocks contained 10 perturbation trials and 40 unperturbed trials per instructed Preparation Time and target size.

Each session began and ended with 2 blocks of free Reaction Time trials. The first two experimental blocks were preceded by a practice block and followed by one practice and 10 experimental blocks of forced Preparation Time trials. Short breaks were inserted after the first, the fifth and the ninth experimental block of forced Preparation Time trials to allow the participants to stand up, stretch their arms and legs, and to drink if necessary.

### Data processing

Data processing was based on the data processing used in previous papers (3, 4). Our primary outcome was the force applied against the perturbation when the hand was 13cm away from the starting position (perturbation trials only). This outcome was obtained from the low-pass filtered force profiles (second-order Butterworth filter with cutoff: 50 Hz). For right perturbation trials, the perpendicular force was sign-reversed in order to pool the data across perturbation direction. For unperturbed trials, we measure the horizontal position of the hand with respect to the straight trajectory connecting the start position to the center of the target. This position is measured when the movement extent was 13cm. All measures were taken at 13cm to ensure that trials where participants anticipatively stopped before the target could be included. A strong correlation (R > 0.8) between the lateral deviation at 13cm and at the end of the movement has previously been reported (4). In an exploratory investigation of how early in the movements the reported effects were present, values of the force were also measured 2cm, 4cm and 5cm into the movement.

The time of movement onset was measured in both perturbed and unperturbed trials based on a velocity threshold of 2cm/s. In both forced Preparation Time and free Reaction Time trials, the reaction time corresponds to the time between the appearance of the target and the start of the movement.

Data processing was conducted in Matlab (readKinTiPR.m, AnalyzeTiPR.m and ExtractTiPR.m on the OSF page) to process the raw data and extract the relevant parameters. Further data processing steps were conducted in Python (Beep_v1.py on the OSF page) to prepare the data for statistical analyses.

### Data Analysis

Statistical analysis was carried out in R. R scripts to reproduce the statistical analysis are available on the OSF page: doi: 10.17605/OSF.IO/8YAHE

### Analysis 1

This analysis aimed to compare the application of the minimum intervention principle in forced Preparation Time and free Reaction Time trials. We hypothesized that participants would make more adjustments when reaching to smaller target apertures compared to larger apertures. Reaches to the smaller target would therefore show more evidence of intervention (i.e. have lower lateral deviations and higher applied forces) compared to reaches made to the larger target. We predicted that this effect of target aperture would be present for both forced Preparation Time trials and free Reaction Time trials. Consequently, three outcomes were examined: 1) the mean and 2) standard deviation of the lateral deviation, and 3) the mean force applied against the perturbation. These outcomes were submitted to a 2×2 ANOVA with the factors condition (forced Preparation Time vs free Reaction Time) and target size (narrow vs wide) as within-subject factors. The aov_car function from the afex package (25) was used to fit the ANOVA model. The implemented analysis can be found in the file FreevsForced_all.R on OSF (doi: 10.17605/OSF.IO/8YAHE)

### Analysis 2

A second analysis examined the effect of preparation time on movement planning. We hypothesized that either the time to compute an optimal movement would be negligible (in which case we would see a stepwise effect) or based on the available preparation time (in which case we would see a gradual increase in changes in the control policy over time). Analyses were conducted examining the variables of 1) mean lateral deviation, 2) standard deviation of the lateral deviation, and 3) mean force applied against the perturbation. These data were submitted to separate 2×5 repeated measures ANOVAs with the factors of Target Size (narrow vs wide) and the available Preparation Time, pooled across five intervals ([-1000ms,50ms], [50ms,200ms], [200ms,350ms], [350ms,500ms] and [500ms,650ms]). Greenhouse-Geisser correction was applied because the sphericity assumption was violated (Degrees of freedom were then multiplied by GG epsilon). Although we recognize that categorizing a continuous variable is not optimal, this was necessary to be able to look at the evolution of the standard deviation in function of Preparation Time. For the analysis of the force outcome, 6 participants were not taken into account because they did not have datapoints in all Preparation Time categories. Analysis 3 deals with the preparation time as a continuous factor for the force outcome. The implemented analysis can be found in the file AnalysisRTSub.R on OSF (doi: 10.17605/OSF.IO/8YAHE)

### Analysis 3

To investigate the influence of reaction on the force applied against the perturbation without discretizing the preparation time variable, we used a linear mixed model with Force as dependent factor and target size as fixed effect and reaction as continuous random effect. Because the influence of the preparation time on force could be subject-dependent, we also added a random slope for the effect of preparation time. This gave us the following model: Force ∼ Target Size*Preparation Time + (Target Size*Preparation Time|Participant). Preparation Time was z-normalized before being entered into the mixed model using the scale() function from R. This linear mixed model was fitted using the “lme4” package of R (26). The implemented analysis can be found in the file MixModelForce_all.R on OSF (doi: 10.17605/OSF.IO/8YAHE)

### Mean comparison

Throughout the paper, mean comparisons were performed with paired t-tests. The results of these t-tests are presented in the figures. To account for the fact that we used two (Fig.4) or five comparisons (Fig.5 and 7) per analysis, we set the significance threshold to 0.01 for these analyses.

### Data availability statement

The pre-processed data (position, velocity and force over time for each trial and each participant) and the processed data (lateral deviation at 13cm, force at 13cm and reaction time for each trial and each participant) are available on the OSF page (doi: 10.17605/OSF.IO/8YAHE). Raw data are available upon request from the first author.

## Results

In this experiment, we asked participants to reach to either a narrow or wide target while either asking them to initiate their movement as quickly as possible (free Reaction Time condition) or imposing a Preparation Time by requiring them to initiate their movement in synchrony with the last beep of a series of four beeps (forced Preparation Time condition). As illustrated in Fig,2, the participants were able to time their movements in synchrony with the last beep although they tended to slightly lead the go-cue by around 40ms for short preparation time to around 140ms for longer instructed Preparation Time. Therefore, participants tended to start slightly earlier than the instructed Preparation Time, which is consistent with previous studies (27, 28). Across all trials, participants were able to time their movement to the go cue. For the short preparation times (<330ms), the movement was initiated less than 100ms before or after the go cue in 70% of the trials or more (from 68% for Preparation Time=1ms to 92% for Preparation Time=264ms). Given the earlier movement initiation discussed above, the adherence plummeted for longer instructed Preparation Times, averaging around 20% (from 25% for instructed Preparation Time of 396ms to 15% for instructed Preparation Time of 594ms). Note that this does not influence the results presented below as we took the actual preparation time (i.e. time between stimulus presentation and the initiation of movement) instead of the instructed one (time between stimulus presentation and final tone). As can be seen on Fig.2, the pattern is very similar for both perturbed and unperturbed trials.

Before looking at the effect of preparation time on our outcomes, we wanted to make sure that the overall behavior was broadly consistent with the minimum intervention principle. Following this principle, participants should have lower error (i.e. have lower lateral deviation of the cursor) for a narrow target than for a wide target and the variability of this error will be larger for wide than for narrow targets (reflected by a lower standard deviation of the lateral deviation of the cursor). In addition, it predicts that the force applied against a perturbation will be larger for narrow than for wide targets.

**Fig. 2:**
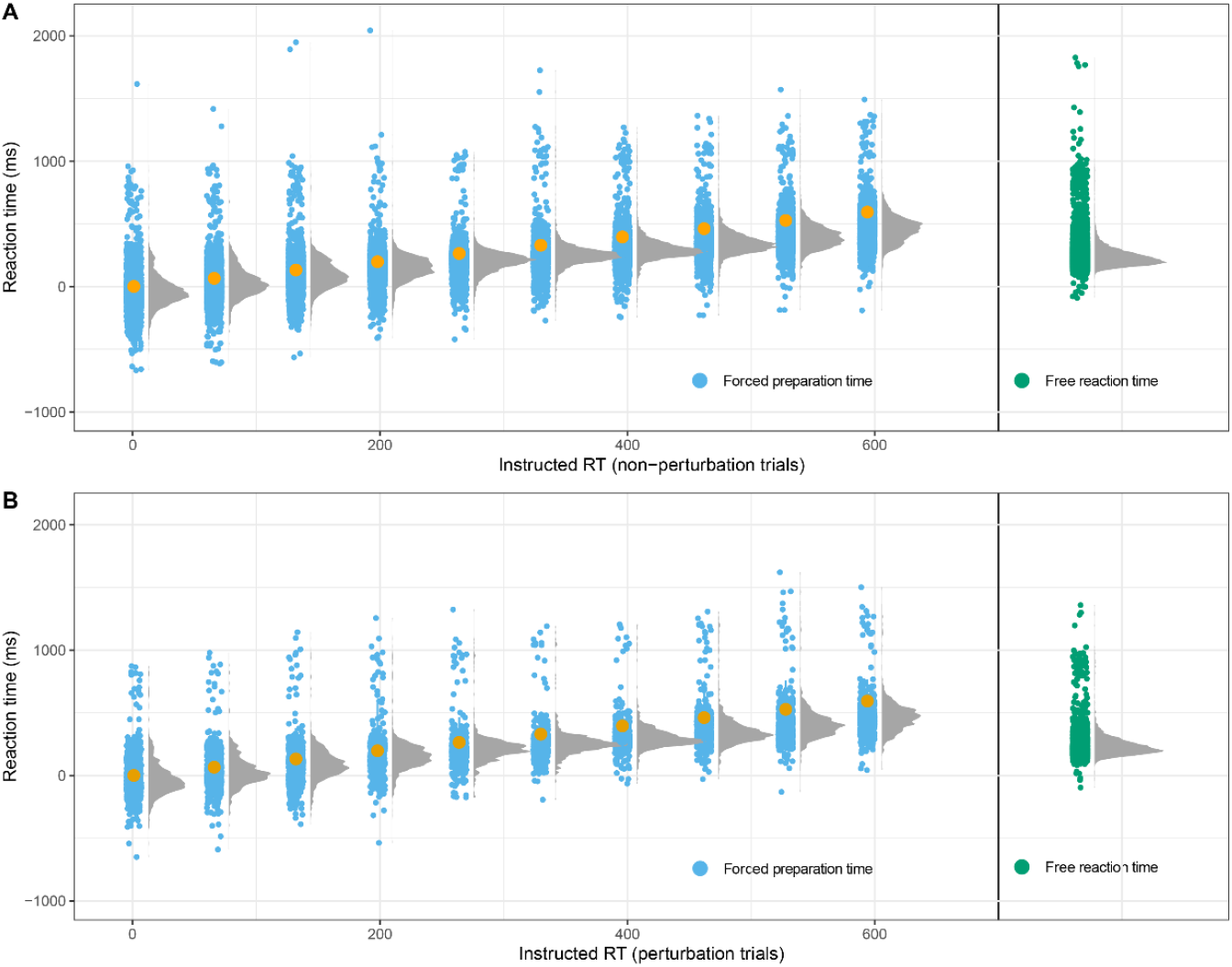
distribution of preparation time of each instructed preparation time for both unperturbed (panel A) and perturbed (panel B) trials. Blue dots correspond to preparation time in the forced Preparation Time condition for the different instructed preparation times (orange dots, between 1 and 594ms by steps of 66ms). Green dots represent the trials in the free Reaction Time condition. Grey histograms represent the preparation time distribution for each instructed preparation time and for the free Reaction Time condition.

These predictions were confirmed in the participants presented in Fig.3. In unperturbed trials (Fig.3, panels A-C), this participant guided his/her hand through the aperture in a swift reaching movements. The inter-trial variability is clearly larger for the wide aperture (Fig.3B) than for the narrower one (Fig.3A). Given that the participants had to move through the aperture within a defined time interval, the hand velocity was close to its maximum when the hand reached the aperture (Fig.3C). In some trials, a perturbation diverted the hand away from the aperture (2cm on the left or right). This participant exerted a gradual increase in force for both the narrow and wide targets and the force was larger for the narrow target than for the wide one (Fig.3D).

**Fig. 3:**
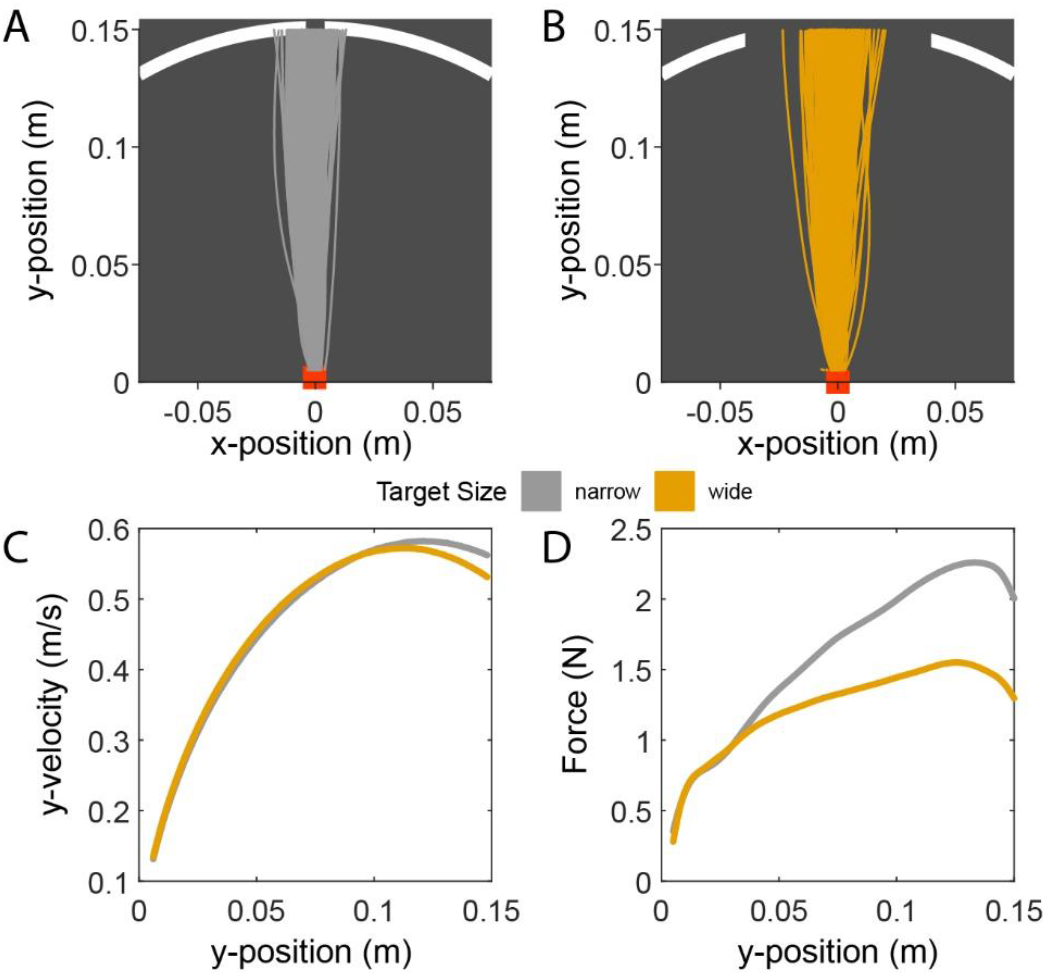
Illustration of the behavior from one participant. Participants were instructed to pass their hand through the aperture of a circle. The aperture could be either narrow (panel A, 1cm) or wide (panel B, 8cm). Panels A and B represent the hand trajectory during the unperturbed trials from the first two free-RT blocks. Panel C represents the velocity of the hand in the same unperturbed trials for the narrow (grey trace) and wide targets (orange trace). Panel D represents the force exerted by the participant against the perturbation.

We measured the different outcomes (position error, variability and force) 13cm into the movement as in our previous publications (3, 4). In the free Reaction Time condition, when movement preparation time was unconstrained, this distance was travelled in approximately 400ms (perturbed trials: 396±4ms, unperturbed trials: 384±4ms, mean ± SE). In the forced condition, these timing remained unchanged (perturbed trials: 408±5ms, unperturbed trials: 398±5ms, mean ± SE). Fig. 4A shows that the force was indeed larger for the narrow target than for the wide target (Analysis 1, main effect of Target size: F(1,39)=84.25, p<0.001). The force applied in the free Reaction Time condition was also larger than in the forced Preparation Time condition (main effect of condition: F(1,39)=11.02, p=0.002). Even though the average force across target was larger in the free Reaction Time condition than in the forced Preparation Time condition, we did not find evidence that this influenced the modulation of force by the target size (interaction between target size and condition: F(1,39)=0.003, p=0.96). Importantly, we found a modulation of force by target size both for the free Reaction Time condition (Fig.4B) and the forced Preparation Time condition (Fig.4C) independently. In short, the behavior of the participants followed the minimum intervention principle in perturbed trials.

**Fig. 4:**
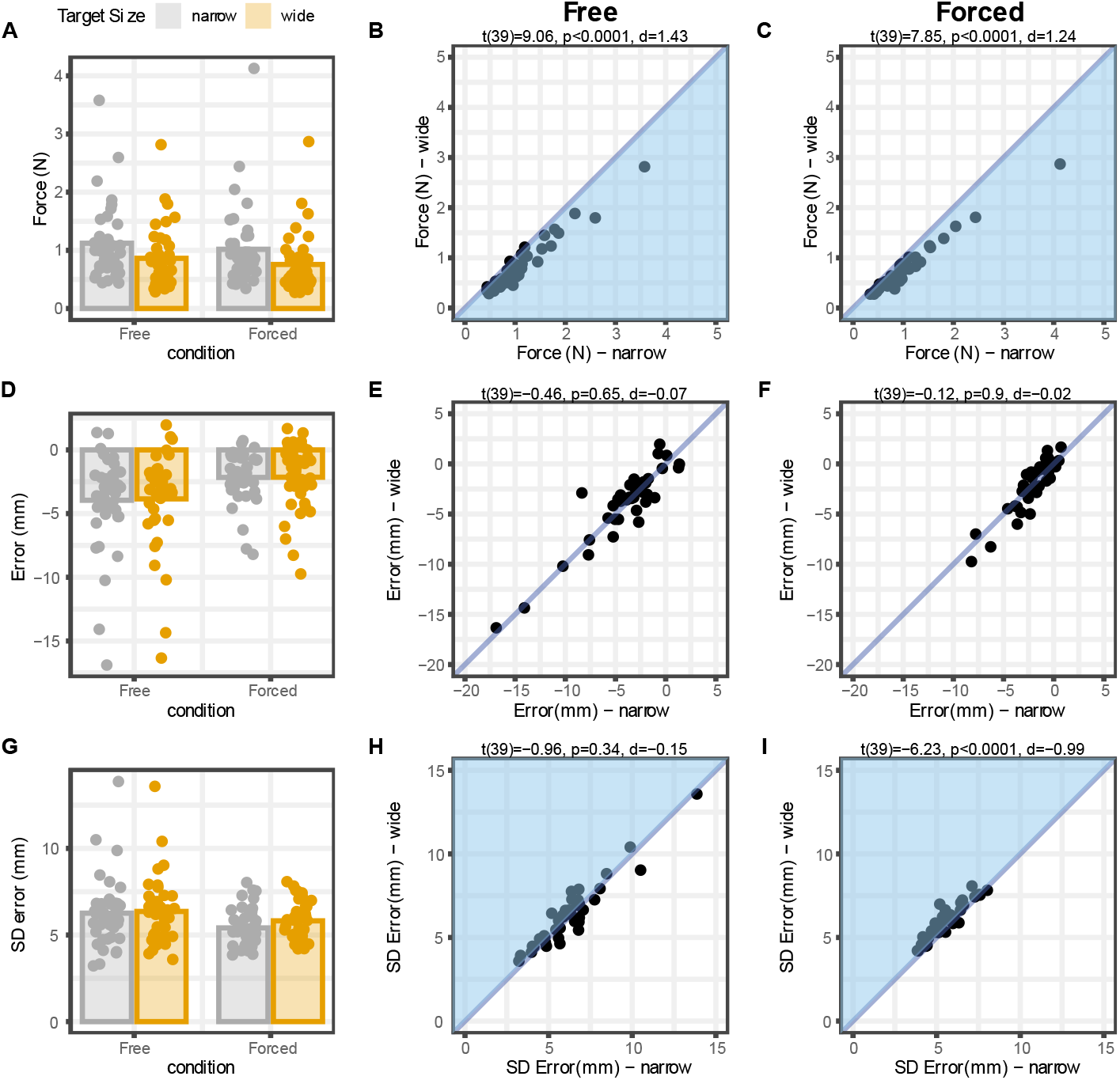
comparison of the free Reaction Time (RT) and forced Preparation Time (PT) condition. First, second and third rows illustrate the force applied in perturbed trials, the average lateral deviation in unperturbed trial and the standard deviation (SD) of the lateral deviation in unperturbed trials, respectively. In the first column, the average outcome is represented for each target size and each condition. Individual datapoints represent the individual datapoints of each participant. Second and third column represent the average value of the outcomes for the narrow target (horizontal axis) vs the wide target (vertical axis) in the free Reaction Time and forced Preparation Time conditions, respectively. Each datapoint represents the average of the outcome for an individual participant. For the force, points under the diagonal are consistent with the minimum intervention principle. For the standard deviation of the error, individual points above the diagonal are consistent with this principle. Statistics above the panels reflect the outcome of paired t-test between the values for the wide target vs the values for the narrow target. Blue areas correspond to areas compatible with the minimum intervention principle.

In unperturbed trials, the results were mixed. We did not find evidence that the accuracy of the reach movement (reflected by the position error; measured as the distance between the center of the target and the final position of the participant’s hand/cursor) was modulated by target size (Fig.4D-F, Analysis 1: main effect of target size: F(1,39)=0.13, p=0.72). Unexpectedly, participants were more accurate in the forced Preparation Time condition than in the free Reaction Time condition (main effect of condition: F(1,39)=25.78, p<0.001, mean+stdev for Free response condition = -0.39+-0.36cm, Forced response condition –0.22+-0,23cm), but the difference in accuracy was actually small and represents a difference of less than 2mm. This behavior was not different for the narrow or wide target (interaction between target size and condition: F(1,39)=0.17, p=0.68). This suggests that the position error cannot reflect the implementation of the minimum intervention principle and will not be investigated further. In contrast, we found that the variability of the reach endpoint error was modulated by target size (Fig.4G, Analysis 1: main effect of target size: F(1,39)=13.83, p=0.0006) and by the condition (main effect of condition: F(1,39)=11.09, p=0.002). Interestingly, the modulation of the variability of reach endpoint by target size was more pronounced in the forced Preparation Time condition (Fig.4I) than in the free Reaction Time condition (Fig.4H, interaction between target size and condition: F(1,39)=8.89, p=0.005). Further analysis demonstrate that, in the forced condition, the variability of the reach endpoint error was larger for the wide target than for the narrow target (t(39)=-6.23, p<0.001, d=-0.99). We did not find evidence that this was the case in the free Reaction Time condition (t(39)=-0.96, p=0.34, d=-0.15).

### Influence of preparation time on the optimality of movements

Our key question is whether the amount of preparation time that is given to the participants will influence the optimality of their movements. If preparation time influences this optimality, we expect that 1) the difference in force between the narrow and wide targets will increase with longer preparation time and that 2) the difference in variability of reach endpoint error between the narrow and wide targets will also increase with preparation time.

Our first approach was to categorize the preparation time of the individual trials into five different categories and to see whether the optimality of the movement was influenced by these categories (Fig.5). We did not find any evidence that the preparation time influenced the modulation of the force applied for the two different target sizes (Analysis 2: interaction between target size and preparation time category: F(2.81,92.81)=0.89, p=0.44) even though there was a clear modulation of the force by target size (main effect of target size: F(1,33)=48.95, p<0.001). As shown on Fig.5, more force was applied for the narrow target than for the wide target for each preparation time category. Effect size (Cohen’s d) of the effect of target size on the force were large and ranged between 0.67 and 1.14. Interestingly, this effect was also true for movement starting less than 50ms before the target appeared (Fig. 5B). In those movements, the movement was initiated before the visual feedback about the target size could influence their control policy. Yet, by the end of the movements, those movements were optimal.

**Fig. 5:**
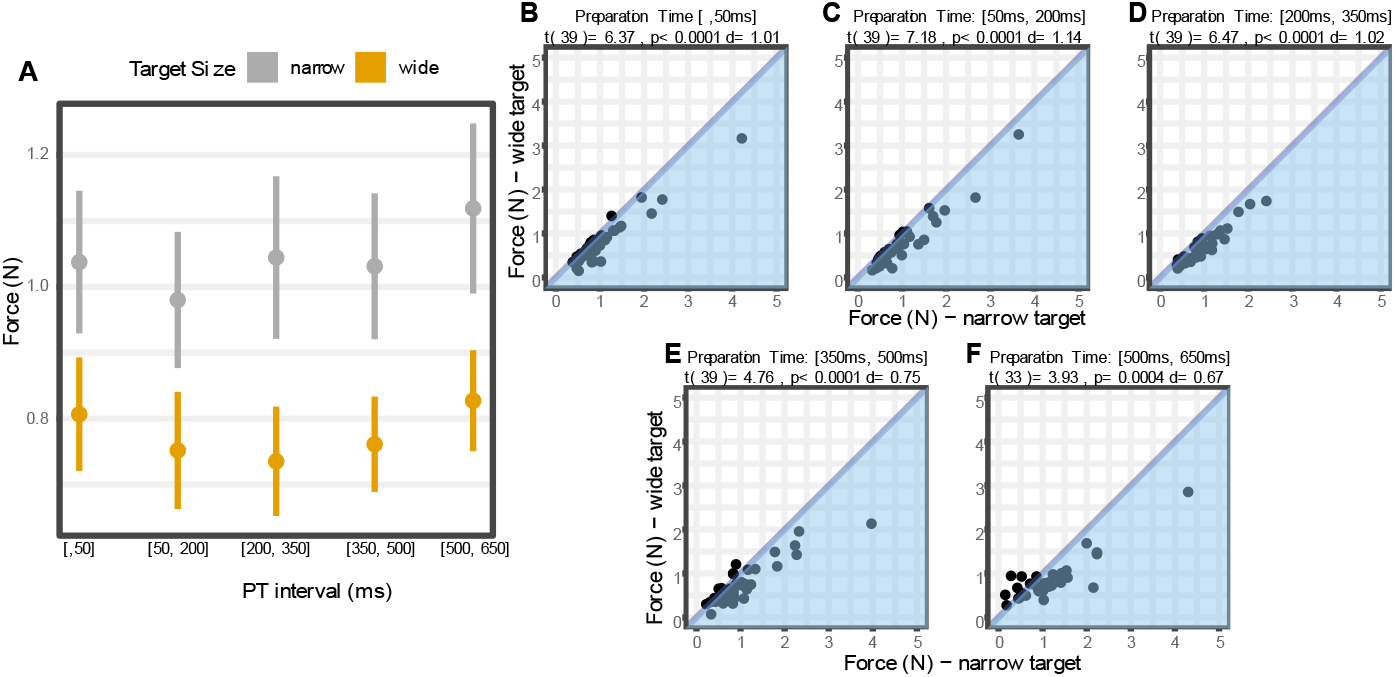
A) Force exerted against the perturbation for the narrow and wide target for each instructed preparation time bin. Each dot represents the mean force for that target width and the error bar represents the standard error of the mean. B) Force exerted against the perturbation for the wide target vs. for the narrow target. Each dot represents the data of an individual participant. Blue areas correspond to areas compatible with the minimum intervention principle.

To take the full spectrum of preparation time into account and to avoid losing power due to the categorization of a continuous variable, we ran a mixed model analysis on the force data (Fig.6, Analysis 3). As expected, the mixed model revealed a strong effect of target size on the force applied against the perturbation (coefficient of the effect of target size: t(38.5)=-7.75, p<0.001) confirming earlier analyses. Again, there was no evidence that preparation time influenced the modulation of force by target size (coefficient of the interaction between target size and Preparation Time: t(336)=-0.307, p=0.76). Upon close inspection of Fig.6, there seems to be a small interaction between Preparation Time and target size whereby the two regression lines appear to diverge with preparation time. Yet, this effect is too small to be detected despite the 40 participants and their 200 perturbation trials each.

**Fig. 6:**
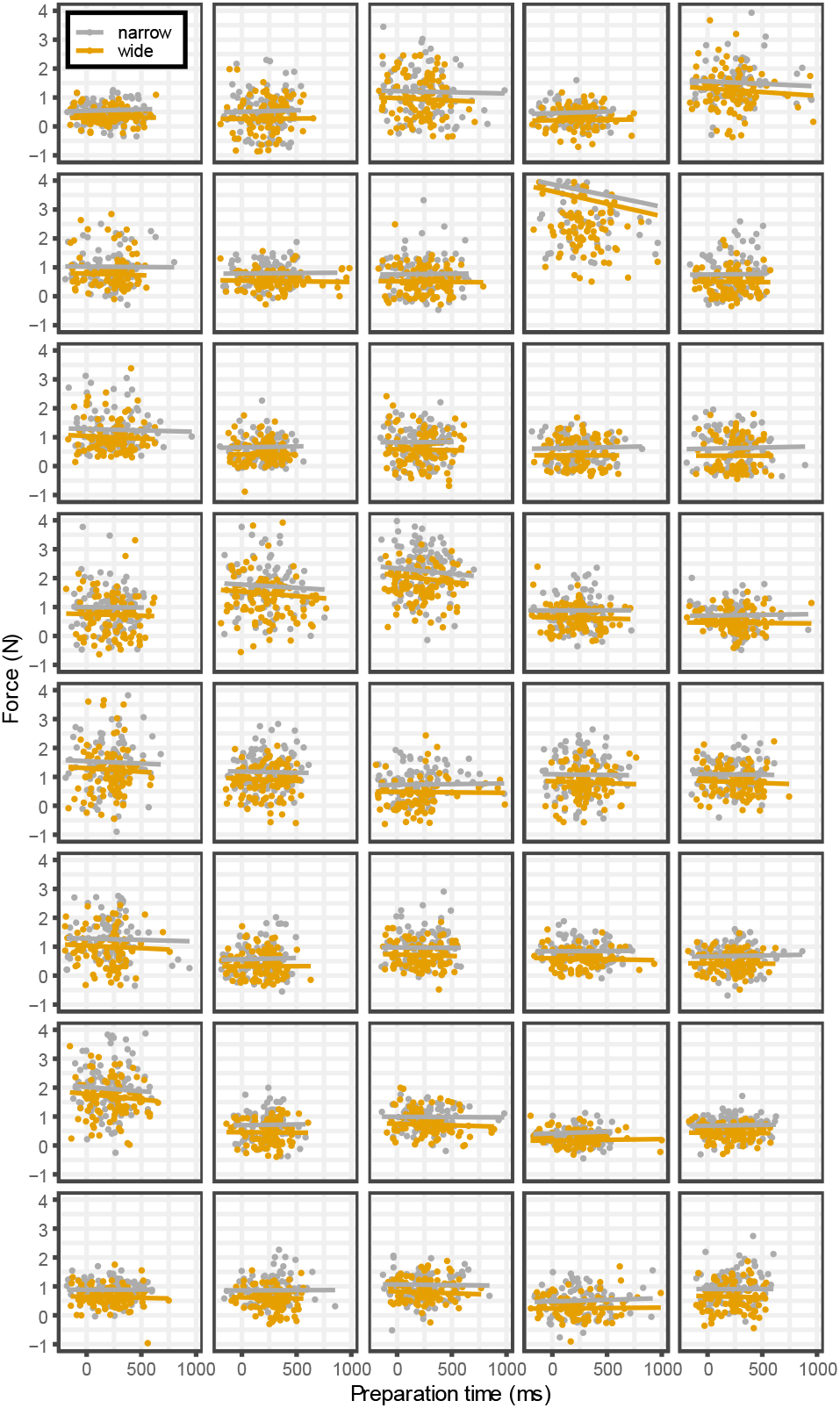
Analysis of data from the Forced Response Condition showing the prediction from the linear mixed model (regression lines) superimposed on the force (N) data for each individual trial in function of preparation time (ms). Each panel represents the data of a single participant (note only one participant of the fourty tested appeared to have a bad individual fit; see column 4, row 2). Each dot represents a single perturbation trial.

Qualitatively similar results were obtained for the standard deviation of the reach endpoint error (Fig.7). This outcome was modulated by target size (Analysis 2: main effect of target size: F(1,39)=19.16, p<0.0001), and we did not find evidence that this modulation differed across preparation time categories (interaction between target size and preparation time categories: F(2.13,83.39)=1.31, p=0.27). In contrast to our results with the force, the effect sizes were much smaller, ranging from 0.14 to 0.82. In addition, the modulation of the variability of reach endpoint error with target size did not reach significance for the early (earlier than 50ms) and late preparation time categories (later than 500ms). Those are also the categories with the least number of trials per participant, making the estimation of variability more difficult.

**Fig. 7:**
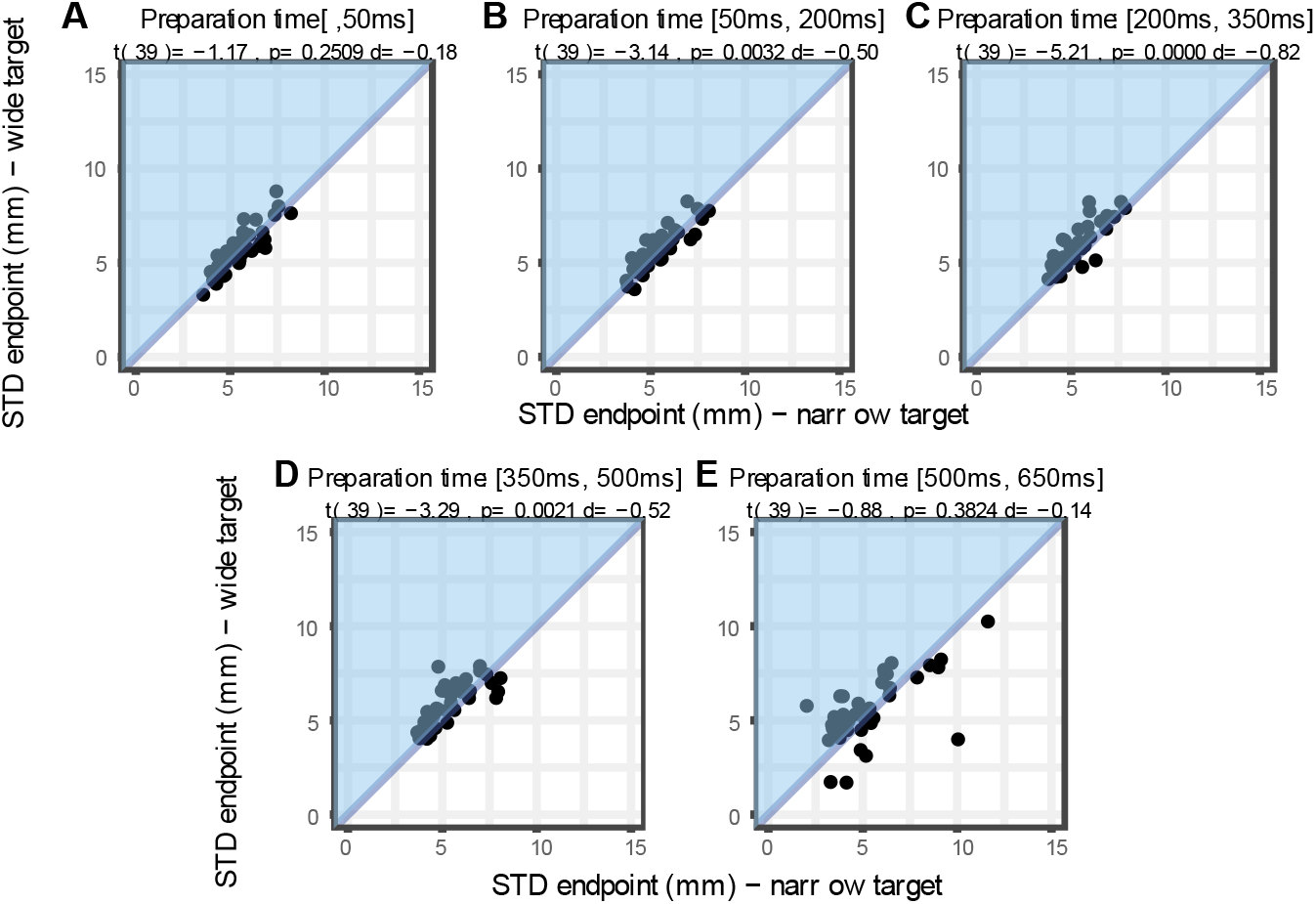
<u>Analysis of data from the Forced Response Condition showing the</u> standard deviation of the lateral position of the end at 13cm for the wide target vs. for the narrow target. Each dot represents the data of an individual participant. Blue areas correspond to areas compatible with the minimum intervention principle (more variability for the wider target).

In the outcomes reported up to now, force was measured when the hand had travelled 13cm. This measure of performance occurs around 400ms after movement onset, leaving ample time for online tuning. Yet, measuring the force earlier in the movement further supports the finding that target size, but not preparation time, influences the force applied against the perturbations. Indeed, measuring it earlier (5cm) into the movement (around 230ms after movement onset) does not change any of the results. However, at that time, the size of the perturbation is much smaller, reducing the signal to noise ratio. Focusing on the force outcome measured at 5cm, we found that the modulation of the force by target size was absent in the Free Reaction Time condition (t(39)= −0.60, p= 0.5550 d= −0.09) but still present in the forced Preparation Time condition (t(39)= 3.53, p= 0.0011 d= 0.56), yielding an interaction between target size and condition (F(1,39)= 5.64, p= 0.023). This absence of difference in the free Reaction Time condition is potentially due to the decreased signal-to-noise ratio in that condition and to the lower number of trials in free Reaction Time compared to forced Preparation Time.

As for the force measured at 13cm, we submitted the force values computed 5cm into the movement from forced Preparation Time condition to an ANOVA (analysis 2) and to the linear mixed model in order to look at the effect of preparation time onto the force. These analyses yielded qualitatively similar results as at 13cm in the forced Preparation Time condition despite the time in the movement being much shorter. When preparation time were separated into different categories (analysis 2), we found that target size influenced the force applied by the participants against the perturbation (main effect of target size: F(1,33)=9.18, p=0.005) but did not find evidence that this difference was modulated by preparation time (interaction between target size and Preparation Time: F(4,132)=0.34, p=0.85). Similarly, in the mixed model (analysis 3), target size modulated force at 5cm (t(313.5)=-3.46, p=0.0006). We did not find any evidence that preparation time influenced the difference in force between the two target widths (t(3118)=-0.245, p=0.81), again consistent with the results at 13cm.

These analyses are limited as it is not possible for us to determine at what point and response to the perturbation would begin to be implemented, and this could vary trial-by-trial. However, we did not find any evidence that target width influenced force outcomes when they were measured at 2 or 4cm in the forced Preparation Time condition (different in force applied between the narrow and wide target measured at 2cm into the movement: t(39)= −1.24, p= 0.22 d=−0.2; measured at 4cm into the movement: t(39)= 0.88, p= 0.38 d= 0.14).

## Discussion

In the present study participants were asked to reach for either a narrow or wide target, initiating their movement as quickly as volitionally possible (free Reaction Time condition), or in a manner that allowed us to experimentally manipulate the time they had available to prepare their movement (forced Preparation Time condition). Across both conditions, when participants experienced a perturbation, their behavior generally followed the minimum intervention principle; they applied greater forces and were more accurate for narrow targets, while the variability of their endpoint error was greater for wide targets.

In contrast to our expectation, the modulation of force with target width did not change in function of the amount of time participants were given to prepare their movements in the forced Preparation Time condition. Strikingly, even when the movement started *before* the information on target width was presented, participants were able to modulate the force applied against the perturbation as well as in the when preparation time was unconstrained. That is, we did not observe any difference in the adaptation of the control policy to task constraints when the control policy was tuned during movement planning or during ongoing movement. Together, these results suggest that the tuning of the control policy is instantaneous and can occur both during movement preparation and execution. Note that the tuning of the control policy could take different forms, and we are agnostic to which form is being implemented. Adaptation to a change in task demands could be implemented by a tuning of the feedback gains of the control policy (19, 29), a change in impedance control (using muscle co-contraction for instance, 30, 31) or a combination thereof.

### How long does it take to tune the control policy?

Our present results are consistent with the proposal that the tuning of the control policy to task demands does not impact reaction time; while participants had more time available to prepare their movements in forced Preparation Time trials with long preparation times, the movements they performed did not differ significantly from those performed with shorter preparation times. This is broadly consistent with previous theories of movement preparation (1), which suggest that movement preparation does not affect reaction time.

Yet, this does not mean that the tuning of the control policy is instantaneous. It is possible that the movement is being initiated while the tuning of the control policy is not fully finished but that this can occur during ongoing movement. A modelling effort on understanding movement preparation suggests that this can be the case (32). In addition, previous studies have demonstrated that it is possible to adapt the tuning of the control policy during ongoing movement (16–18), yet is currently unclear how much time this takes. De Comite and colleagues (17) showed that the tuning of the control policy representing the adaptation of the control policy to the task demands was completed within 150ms. Given visuomotor delays, they estimated that it takes 50ms to update the control policy during ongoing movement. Yet, the timing was not systematically investigated in this task. In a postural task, Yang and colleagues systematically vary the timing of the presentation of task instruction and found that long-latency stretch reflexes can reflect the right instruction within 90ms of its presentation (33). Our data does not allow us to estimate the amount of time necessary to adjust the control policy more accurately than in the previous studies. Indeed, we do not know when the perturbation was detected by the brain (or whether it was detected at all) as the size of the perturbation increases gradually with the distance travelled by the hand. If we hypothesize that it takes 50-100ms for the brain to detect the perturbation, that on average people initiate their movements at the time of the go cue, and that sensorimotor delays amount 100ms, the observed response at 5cm (around 230ms after movement onset) would be consistent with a delay of 80-100ms to adjust the control policy to task demands. Yet, this experiment is not suited to answer this question. Therefore, we can’t be sure of this estimate and future studies based on EMG should establish this time more precisely.

### Why is this at all surprising?

Both the theory (1) and previous research (17) suggested that it took little time to adjust the control policy to the control policy. The framework of Wong et al. (1) even predicted that it would not affect reaction time. Yet, we were willing to confirm this experimentally because of the existence of a switching cost for switching between two control policies (4). When wide or narrow targets are presented for many trials in a row, the force exerted against a perturbation is more influenced by target width than when target width varies from trial to trial (3, 4). This suggests that adapting the control policy to task demands is an effortful process, yet this effortful process does not impact reaction time as is typical for switching cost in the cognitive domain (5).

### Does motor planning actually exist?

A surprising result of our study is that there was absolutely no advantage in tuning the control policy before movement onset compared to during movement execution. That is, the tuning of the control policy was similar for trials where the movement started after the presentation of the target and for those where the movement was initiated before the visual information about target width was displayed. This is consistent with the idea that there can be a large overlap between planning and execution (34, 35) and that movements can be initiated before movement preparation is finished (22). One can also prepare the next movement during ongoing movement (36). Yet, the present result and the aforementioned studies ask the question whether the computation of the -optimal-motor control policy is actually part of the motor planning stage, or whether this embedding results from the use of delayed reaching task paradigms where movement planning and execution are artificially separated. Indeed comparing delayed to quasi-automatic reaches, Lara and colleagues (34) suggested that the motor preparatory stage was present for both types of trials but that the motor preparatory stage was taking very little time in quasi-automatic reaches. They estimated that the motor preparatory stage could take as little as 40ms, which would be undetectable in the present experiment. In summary, the present study adds to a series of recent results that demonstrate that the separation between motor planning and motor execution is not as discrete as was once thought, and that the motor preparation stage can happen during movement.

### How does this view fit with motor preparation in the brain?

In the motor cortex, preparation and execution of movements are well segregated at the neural levels. In the dynamical system framework (37), motor preparation results in setting the population state at a given point reflecting the initial conditions (38) that is followed by rotational dynamics linked to movement execution (39). In addition, neural activity linked to movement preparation and to execution lives in orthogonal subspaces (40). In the idea that movements can be initiated before motor preparation is fully completed, it has been reported and theorized that different endpoints in movement preparatory activity can be reached when movement are initiated without loss of movement accuracy (32, 41). Even with quasi-automatic movements, the motor preparation stage takes place but partially overlap with movement execution (34). Yet, it is unclear how modulation of the control policy by task demands will influence these mechanisms. Would a change in the control policy be reflected in different neural activity in preparatory dimensions, or in different rotational dynamics/movement execution dimensions?

The efficacy of measurements of performance in forced Preparation Time conditions is critically dependent upon participant’s adherence to task instructions. In the present experiment, participants were generally able to time their movements in synchrony with the last tone, but tended to start slightly earlier than the instructed time with the forced Preparation Time was longer. Informal examination of prior experiments indicates that participants are more likely to delay the initiation of their movements when a very short time to prepare an action is imposed (Hardwick et al., 2019, Vleugels et al., 2020).

In conclusion, the present study indicates that increasing the amount of time available for movement preparation does not lead to significant differences in movement planning. This is consistent with previous work which suggests that motor plans can be generated almost instantaneously. Moreover, our results indicate that movements prepared volitionally (in free Reaction Time conditions) and with limited time (in forced Preparation Time conditions) are broadly equivalent, as both are in general accord with the principles of optimal control. These results point to the fact that the frontier between movement planning and execution is not as clear as is often depicted.

## Acknowledgement

We thank Pauline Belloy and Sanne Michielsen for help with data collection.

